# Short- and long-term effects of amoxicillin/clavulanic acid or doxycycline on the gastrointestinal microbiome of growing cats

**DOI:** 10.1101/2021.05.28.446115

**Authors:** Evangelia M. Stavroulaki, Jan S. Suchodolski, Rachel Pilla, Geoffrey T. Fosgate, Chi-Hsuan Sung, Jonathan A. Lidbury, Jörg M. Steiner, Panagiotis G. Xenoulis

## Abstract

Antibiotic treatment in early life influences gastrointestinal (GI) microbial composition and function. In humans, the resultant intestinal dysbiosis is associated with an increased risk for certain diseases later in life. The objective of this study was to determine the temporal effects of antibiotic treatment on the GI microbiome of young cats. Fecal samples were collected from cats randomly allocated to receive either amoxicillin/clavulanic acid (20 mg/kg q12h) for 20 days (AMC group; 15 cats) or doxycycline (10 mg/kg q24h) for 28 days (DOX group;15 cats) as part of the standard treatment of upper respiratory tract infection. In addition, feces were collected from healthy control cats (CON group;15 cats). All cats were approximately two months of age at enrolment. Samples were collected on days 0 (baseline), 20 or 28 (AMC and DOX, respectively; last day of treatment), 60, 120, and 300. DNA was extracted and sequencing of the 16S rRNA gene and qPCR assays were performed. Fecal microbial composition was different on the last day of treatment for AMC cats, and 1 month after the end of antibiotic treatment for DOX cats, compared to CON cats. Species richness was significantly greater in DOX cats compared to CON cats on the last day of treatment. Abundance of Enterobacteriales was increased, and that of Erysipelotrichi was decreased in cats of the AMC group on the last day of treatment compared to CON cats. The abundance of the phylum Proteobacteria was increased in cats of the DOX group on days 60 and 120 compared to cats of the CON group. Only minor differences in abundances between the treatment groups and the control group were present on day 300. Both antibiotics appear to delay the developmental progression of the microbiome, and this effect is more profound during treatment with amoxicillin/clavulanic acid and one month after treatment with doxycycline. Future studies are required to determine if these changes influence microbiome function and whether they have possible effects on disease susceptibility in cats.

## Introduction

Antibiotic discovery represents one of the most important achievements in the history of medicine [1]. However, overuse of antibiotics compromises their health benefits because of the development and dissemination of antibiotic resistant genes. The development of multidrug resistant bacteria is associated with higher morbidity, mortality, and hospitalization costs [2]. Another reason to set boundaries on the extended use of antibiotics is their impact on the gastrointestinal (GI) microbiome [3]. The extent that the microbiome is affected by antibiotics has become apparent after the application of “omics” approaches in research that allow the assessment of whole microbial communities and their functions [4].

The GI microbiome is a community of microorganisms and has been called “a hidden organ” [5]. This community of microorganisms is responsible for maintaining colonization resistance and produces substances with an impact on the host’s metabolism, immune system development and response, and appears to participate in the communication among different organs as well as in the manifestation and progression of diseases [6-14].

The term GI dysbiosis is used to describe the compositional and functional alterations of the GI microbiome in response to exogenous factors and/or the health status of the host [15]. Antibiotic-induced dysbiosis is characterized by a decrease in bacteria beneficial for the host (“health-associated bacteria”), allowing overgrowth of potentially pathogenic bacteria, and a shift in microbially derived metabolic products [15, 16]. Antibiotic-induced microbial shifts can persist long term, and the abundances of some bacterial taxa might never return to their initial state. Other members of the microbiome, including the mycobiome and the virome are also affected by antibiotics, highlighting a global imbalance among members of microorganisms not directly inhibited by antibiotics [17]. Antibiotic-induced dysbiosis depends on the spectrum of antibacterial activity, type, duration, dosage, and route of administration in addition to individual host characteristics [18, 19].

The GI microbiome appears to be more susceptible to antibiotics when administered early in life. During that period, maturation of the immune system takes place concurrently with microbiome maturation. Antibiotics result in exposure of the host to a reduced number of microbes in the gut, as well as altered microbial signals by the host’s immune system [20]. In addition, antibiotics administered early in life appear to delay the developmental progression of the microbiome into an adult-like state [21, 22]. Previous studies have shown that children exposed to antibiotics were more likely to develop inflammatory bowel disease [23, 24], obesity [25, 26], or asthma [27, 28] during childhood. Currently, limited data is available for cats. In one-study, all cats previously treated with amoxicillin/clavulanic acid and pradofloxacin developed diarrhea after experimental infection with enteropathogenic *E*.*coli* in contrast to non-treated cats, none of which developed clinical signs [29]. This study highlights that similarly to humans, antibiotic-induced dysbiosis likely reduces colonization resistance in cats.

Previous molecular studies investigating the effects of antibiotics on the feline GI microbiome have enrolled healthy laboratory born and bred domestic shorthair cats. In these studies, cats were adults, but belonged to various age groups and were fed the same diet for the duration of each study [30-32]. In humans, antibiotics with an anaerobic spectrum of activity seem to have a more profound and prolonged effect on the gut microbiome, given that 95% of the GI bacteria are anaerobic [33].

Administration of clindamycin affected the feline microbiome and metabolome long-term, with changes persisting for at least 2 years after withdrawal of the antibiotic [31]. Amoxicillin-clavulanic acid is effective mainly against gram positive microorganisms and microbial shifts were still detected 7 days after its withdrawal [30]. No studies to date have investigated the effect of doxycycline and antibiotic treatment in general on the gastrointestinal microbiome of young cats until they reach maturity.

The aim of this study was to describe and compare the fecal microbiome of cats receiving amoxicillin/clavulanic acid or doxycycline and control cats not receiving antibiotics and follow them up over a period of 10 months. A second goal was to describe the normal age-related changes of the feline microbiome changes during development.

## Materials and methods

### Cats

The protocol was reviewed and approved by the Animal Ethics Committee of the University of Thessaly, Greece (AUP number: 54/13.2.2018). A total of 72 eight-week-old rescue domestic shorthair (DSH) cats were enrolled in the study. Forty-four out of 72 cats were diagnosed with upper respiratory tract disease (URTD) before inclusion into the study. Diagnosis was based on a typical clinical presentation, including conjunctivitis, blepharospasm, ocular and/or nasal discharge, nasal congestion, sneezing, and/or coughing. The cats were treated with antibiotics (see Treatment) as part of the standard treatment for this condition. In addition, 26 clinically healthy cats or cats with very mild URTD that did not require antibiotic treatment were enrolled as controls.

Cats were either housed in foster homes or in individual cages at the Clinic of Medicine at the Faculty of Veterinary Science of the University of Thessaly. All cats were eventually adopted into private homes by the end of the study and owners signed an informed owner consent form. Upon initial enrollment, cats were kept under observation for a few days in case they developed clinical signs of GI disease. A physical examination was performed and antiparasitic treatment (Broadline, Boehringer Ingelheim) was administered to each cat before inclusion into the study.

Data including sex, body weight, body condition score (BCS), presence of diarrhea and vomiting, temperature, and heart rate were recorded. Evaluation of BCS and fecal score (FS) was based on previously published scoring systems [34, 35]. Concurrent health conditions were recorded, and cats were excluded if these were severe enough to require hospitalization. All cats were on the same diet (GEMON Cat Breeder Kitten) for the duration of the study, to ensure that differences attributed to diet did not affect the results. No more than two related cats were included in the same group to ensure that relatedness did not impact the results. All cats were vaccinated according to recent vaccination guidelines [36].

### Treatments

Cats with URTD were randomly allocated to receive either amoxicillin/clavulanate at 20 mg/kg q 12 h for 20 days (n=23, AMC group) or doxycycline at 10 mg/kg q 24 h for 28 days (n=21, DOX group). These antibiotics were chosen because they constitute recognized first line treatments for URTD in cats [37]. In addition, 26 clinically healthy cats were enrolled as controls and did not receive any antibiotics during the study period (n=26, CON group).

### Sample collection and follow-up period

Fecal samples were collected from each cat on days: 0 (all groups; one day after initial presentation and antiparasitic treatment), 20 (AMC group; last day of antibiotic treatment for AMC group), 28 (DOX and CON groups, last day of antibiotic treatment for DOX group), 60 (all groups), 120 (all groups), and 300 (all groups). Naturally voided fecal samples were collected from the litter box and placed into Eppendorf tubes. For cats that were adopted, owners were instructed to collect fecal samples from the litter box, freeze them over night and either bring them to the clinic or ship them packed with icepacks by overnight courier. Upon receipt, samples were immediately stored at -80°C pending analysis. On each sampling day, cats underwent a physical examination and the same data as for initial presentation were collected for all cats at all sampling times.

### DNA extraction

Genomic DNA was extracted from 100 mg of each fecal sample using a MoBio PowerSoil® DNA isolation kit (Mo Bio Laboratories, USA) according to the manufacturer’s instructions.

### 16S rRNA sequencing

Illumina sequencing of the bacterial 16S rRNA genes was performed using primers 515F (5’-GTGYCAGCMGCCGCGGTAA) [38] to 806RB (5’-GGACTACNVGGGTWTCTAAT) [39] at the MR DNA laboratory (Shallowater, TX).

Sequences were processed and analyzed using a Quantitative Insights Into Microbial Ecology 2 (QIIME 2) [40] v 2018.6 pipeline. Briefly, the sequences were demultiplexed and the ASV table was created using DADA2 [41]. Prior to downstream analysis, sequences assigned as chloroplast, mitochondria, and low abundance ASVs, containing less than 0.01% of the total reads in the dataset were removed. All samples were rarefied to even sequencing depth, based on the lowest read depth of samples, to 8,275 sequences per sample. The raw sequences were uploaded to NCBI Sequence Read Archive under project number SRP16253.

Alpha diversity was measured with the Chao1 (richness), Shannon diversity (evenness) and observed ASVs (richness) metrics within QIIME2. Beta diversity was evaluated with the weighted and unweighted phylogeny-based UniFrac [42] distance metric and visualized using Principal Coordinate Analysis (PCoA) plots, generated within QIIME2.

### Quantitative PCR (qPCR)

Quantitative PCRs were performed for selected bacterial groups that are commonly altered in canine and feline gastrointestinal disorders: total bacteria, *Faecalibacterium* spp., *Turicibacter* spp., *Streptococcus* spp., *Escherichia coli, Blautia* spp., *Fusobacterium* spp., *Clostridium hiranonis, Bifidobacterium spp*., and *Bacteroides* spp. The qPCR cycling, the oligonucleotide sequences of primers and probes, and respective annealing temperatures for selected bacterial groups have been described previously [43, 44].

### Statistical analysis

Statistical analyses were performed using statistical software packages (SPSS version 23.0; and Prism version 9.0, GraphPad Software). For clinical data, a Kolmogorov-Smirnov test was used to assess the normality assumption. Clinical data did not pass normality testing, and therefore Kruskal-Wallis tests were used for among group comparisons while Friedman tests were used for within group comparisons. Pairwise comparisons were performed using Dunn’s post hoc tests to determine which group categories were significantly different from each other as well as which time points were significantly different.

To determine differences in microbiome composition among and within the study groups, the analysis of similarities (ANOSIM) was performed using the statistical software package PRIMER 7 (PRIMER-E Ltd., Lutton, UK) based on the unweighted and weighted UniFrac distance matrices. Differences in alpha diversity indices and differences in the abundances of bacterial taxa among and within groups were determined using a linear mixed model. Data were rank transformed prior to statistical analyses due to violation of the normality assumptions. Microbial compositions were initially screened for differences among groups with p values adjusted for multiple hypothesis testing using the Benjamini and Hochberg False discovery rate (FDR) and overall significance set at p < 0.05. For comparisons that were significant after FDR adjustment, a linear mixed model was fit including time, group, and the interaction between time and group as fixed effects and cat as a random effect. Multiple pairwise post hoc comparisons were adjusted using Bonferroni correction.

## Results

### Clinical data

Twenty-seven cats were excluded from the study because of owner non-compliance (7/72), death (9/72; 1 due to car accident, 1 due to fall from a balcony, 1 due to feline infectious peritonitis, 1 due to heart failure, while 5 had unknown cause of death), they required a second course of antibiotics (5/72), use of antifungal treatment (3/72), or escape from home (3/72). Fifteen cats in each treatment group (45 cats total) completed the study. These included 25 males and 20 females.

Metagenomic analysis and clinical data assessment were only performed for the cats that completed the study.

On day 0, cats of the AMC group had significantly lower body weights (BW) (median 0.61 kg, range 0.37-0.95 kg) compared to CON cats (median 0.74 kg, range 0.52-1.4 kg) (p =0.026; Table 1). No other BW or BCS differences were identified among groups. On day 0, cats belonging to the DOX group had a significantly higher fecal score (FS) (median 4/7, range 2/7-7/7), i.e., had more commonly abnormal fecal consistency, compared to CON cats (median 2/7, range 1/7-6/7) (p=0.045). On days 20/28 and 60, AMC cats had a significantly higher FS (day 20, median 4/7, range 1/7-6/7; day 60, median 3/7, range 1/7-6/7) compared to CON cats (days 28 and 60, median 2/7, range 1/7-3/7) (p <0.05). Clinical data and p values from the remaining timepoints are listed in Table 1.

**Table 1:** Clinical characteristics of cats included in the study.

|  | Body weight (kg) |  |  |  |  |  | P value |
| --- | --- | --- | --- | --- | --- | --- | --- |
|  | AMC |  | DOX |  | CON |  |  |
|  | Median | Range | Median | Range | Median | Range |  |
| Day 0 | 0.61 | 0.37-0.95 | 0.68 | 0.39-1.20 | 0.74 | 0.52-1.40 | 0.026 |
| Day 20/28 | 1.00 | 0.80-1.56 | 1.18 | 0.87-1.73 | 1.26 | 0.77-1.60 | 0.217 |
| Day 60 | 1.70 | 1.36-1.80 | 1.70 | 1.25-2.00 | 1.92 | 1.15-2.50 | 0.120 |
| Day 120 | 2.50 | 1.98-3.79 | 2.62 | 1.99-3.05 | 2.80 | 1.69-3.80 | 0.746 |
| Day 300 | 4.10 | 2.70-5.75 | 4.19 | 3.33-5.80 | 4.00 | 2.30-6.00 | 0.797 |
|  | Body condition score (1 to 9) |  |  |  |  |  | P value |
|  | AMC |  | DOX |  | CON |  |  |
|  | Median | Range | Median | Range | Median | Range |  |
| Day 0 | 4 | 2-6 | 4 | 3-5 | 4 | 3-5 | 0.107 |
| Day 20/28 | 4 | 4-5 | 4 | 4-5 | 4 | 4-5 | 0.717 |
| Day 60 | 4 | 4-6 | 4 | 4-5 | 4 | 3-5 | 0.651 |
| Day 120 | 4 | 4-6 | 4 | 4-6 | 4 | 4-6 | 0.935 |
| Day 300 | 4 | 3-7 | 5 | 4-6 | 5 | 3-6 | 0.281 |
|  | Fecal score (1 to 7) |  |  |  |  |  | P value |
|  | AMC |  | DOX |  | CON |  |  |
|  | Median | Range | Median | Range | Median | Range |  |
| Day 0 | 3 | 2-6 | 4 | 2-7 | 2 | 1-6 | 0.045 |
| Day 20/28 | 4 | 1-6 | 3 | 2-5 | 2 | 1-3 | <0.001 |
| Day 60 | 3 | 1-6 | 3 | 1-5 | 2 | 1-3 | 0.035 |
| Day 120 | 2 | 1-5 | 2 | 1-5 | 2 | 1-4 | 0.221 |
| Day 300 | 2 | 1-5 | 2 | 1-3 | 2 | 1-3 | 0.195 |

AMC, cats treated with amoxicillin/clavulanic acid for 20 days; DOX, cats treated
with doxycycline for 28 days; CON, healthy cats that did not receive antibiotics.
Bolded p-values indicate a statistically significant difference between groups.

## 1. Effect of aging on the microbiome of untreated cats

### 1.A) Sequence analysis - alpha and beta diversity

High interindividual variations in bacterial abundances were observed in all groups on day 0 and within the CON group significant changes occurred over time. These changes were attributed to the process of microbial maturation, therefore results from this group are discussed separately. In total, the sequence analysis of the 225 fecal samples yielded 1,861,875 quality sequences. There were no differences in any of the species richness and evenness indices over time in control cats (Table 2).

**Table 2:**
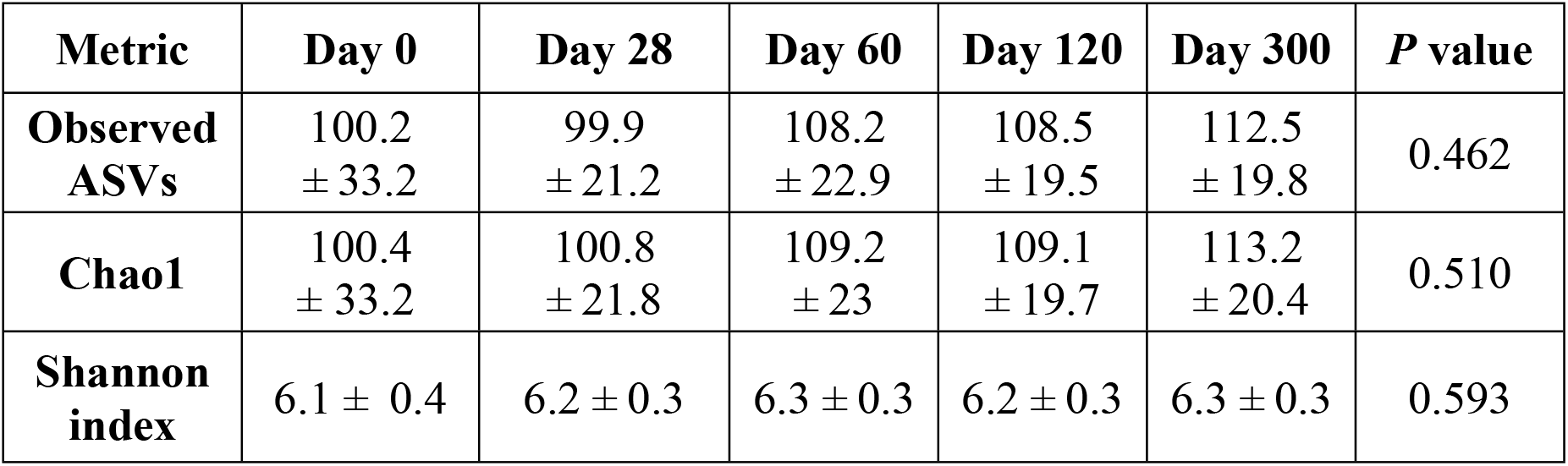
Alpha diversity metrics (mean ± standard deviation) of control cats. There were no significant changes in indices of diversity over time

| Metric | Day 0 | Day 28 | Day 60 | Day 120 | Day 300 | *P* value |
| --- | --- | --- | --- | --- | --- | --- |
| **Observed ASVs** | 100.2<br>± 33.2 | 99.9<br>± 21.2 | 108.2<br>± 22.9 | 108.5<br>± 19.5 | 112.5<br>± 19.8 | 0.462 |
| **Chao1** | 100.4<br>± 33.2 | 100.8<br>± 21.8 | 109.2<br>± 23 | 109.1<br>± 19.7 | 113.2<br>± 20.4 | 0.510 |
| **Shannon index** | 6.1 ± 0.4 | 6.2 ± 0.3 | 6.3 ± 0.3 | 6.2 ± 0.3 | 6.3 ± 0.3 | 0.593 |

However, the phylogenetic community structure clustered significantly different over time (p < 0.05) and was increasingly more distinct as cats were getting older based on the increasing ANOSIM effect size of unweighted and weighted UniFrac distances (Fig 1).

**Fig 1.**
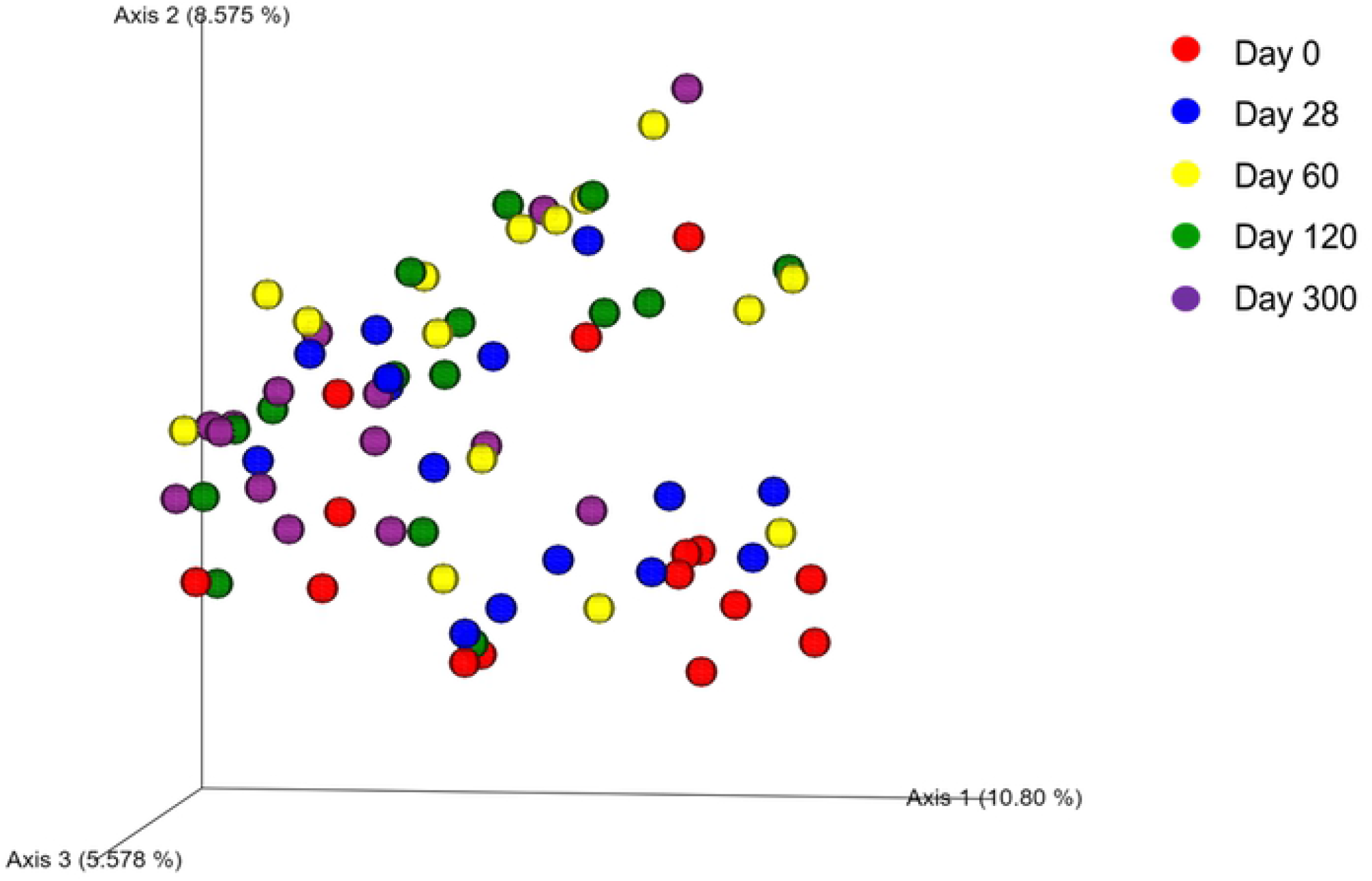
Principal Coordinate Analysis of unweighted UniFrac distances of 16S rRNA genes representing the differences in microbial community composition within the control group on day 0 (red circles), day 28 (blue circles), 60 (yellow circles), 120 (green circles), and 300 (purple circles).

### 1.B) Sequence analysis – abundance of individual bacterial taxa

At 2 months of age (day 0) the most prevalent phylum (regardless of the group) was Firmicutes (63.5%), followed by Actinobacteria (13.9%), Bacteroidetes (11.6%), Proteobacteria (6.0%), and Fusobacteria (4.9%). The abundance of Proteobacteria was significantly reduced to less than 1% (p = 0.009) by 4 months of age in the control cats (Fig 2). Table S1 contains summary statistics for all taxonomic classifications (i.e., phylum, class, order, family, genus, and species).

**Fig 2.**
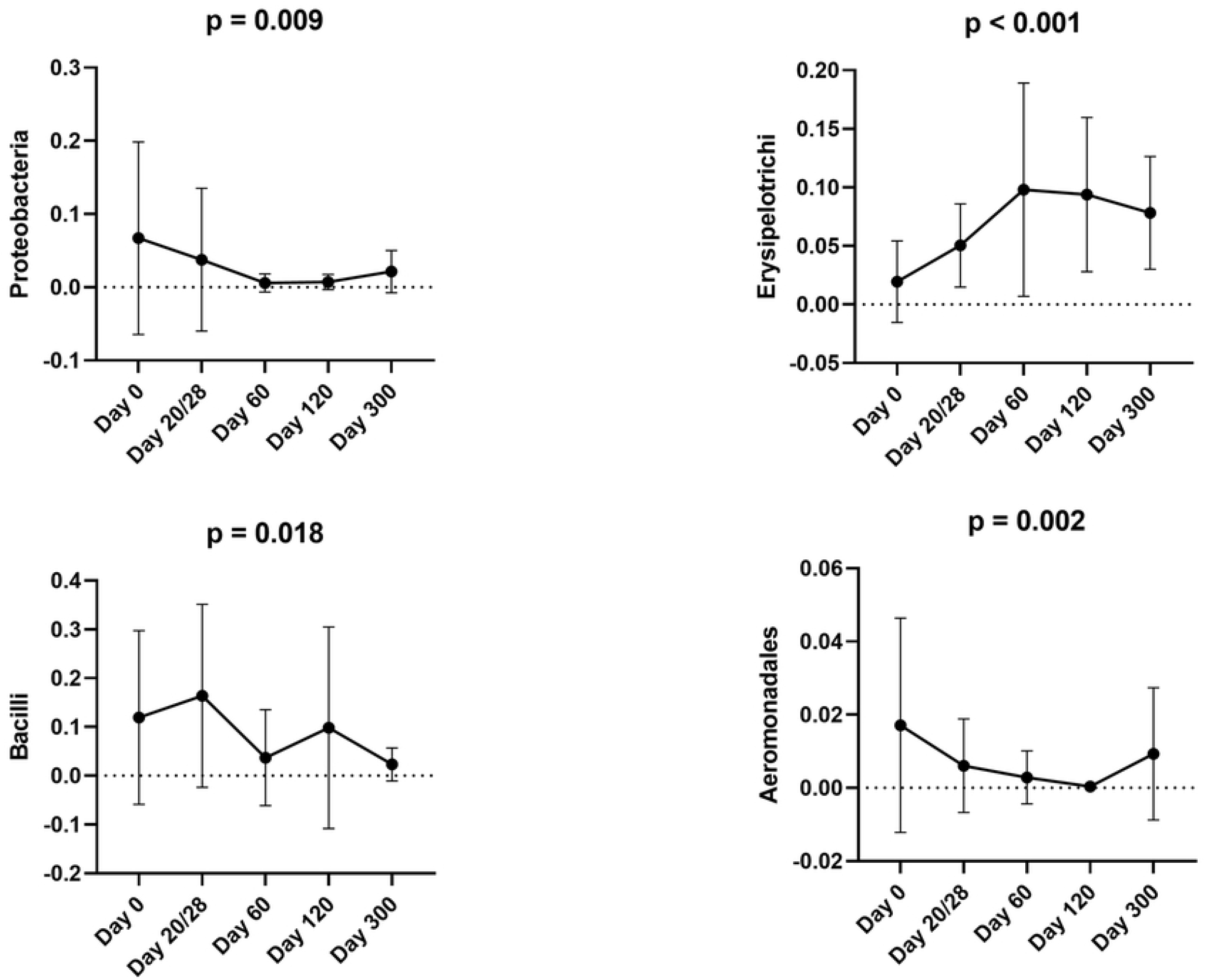
Bacterial groups that significantly changed over time within the control group based on sequence analysis. Means and standard deviations are displayed.

Clostridia, Clostridiales, and Lachnospiraceae, were the most prevalent class, order, and family, respectively, present in fecal samples from control cats during their first year of age. In addition, *Blautia* spp., *Collinsella* spp., *Lactobacillus* spp., *Bifidobacterium* spp., *Bacteroides* spp., and unclassified Lachnospiraceae constituted the predominant genera.

The majority of differences in the abundances of bacteria within the control cats occurred between 2 and 6 months of age. The abundance of Gammaproteobacteria significantly decreased from 5.5% at 2 months to 3.2% at 3 months of age (p = 0.007) and that of Enterobacteriales from 3.7% to less than 0.5% (p = 0.009) during the same period. The abundance of Erysipelotrichi increased from 1.9% at 2 months to 5% at 3 months of age (p = 0.030) (Fig 2). The abundance of Bacilli reduced from 16.4% at 3 months to 3.7% at 4 months of age (p = 0.018). The only changes observed after 6 months of age included an increase in the abundance of Aeromonadales (p = 0.002) (Fig 2).

### 1.C) Quantitative polymerase chain reaction (qPCR) for selected bacterial groups

In the CON group, *E*.*coli* decreased (p < 0.001), and *Faecalibacterium* spp. increased (p = 0.032) from 2 to 3 months of age (Fig 3). Table S2 contains a summary of all bacterial taxa analyzed by qPCR.

**Fig 3.**
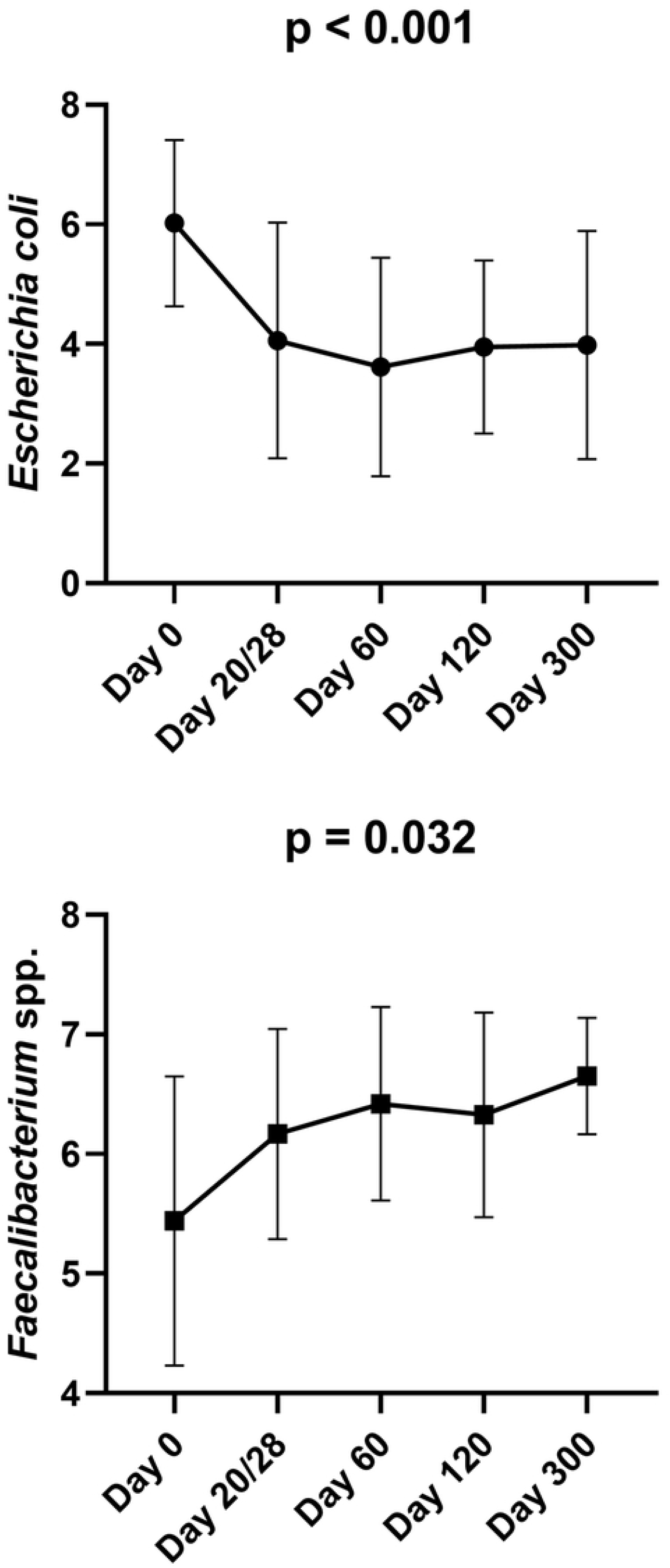
Bacterial groups that significantly changed over time within the control group based on qPCR analysis. Means and standard deviations are displayed.

## 2. Effect of antibiotics on the GI microbiome

The effect of antibiotics on the GI microbiota was assessed based on comparisons among groups on the same timepoints. Because the GI microbiome normally changes over time as it evolves towards maturity, overtime comparisons withing the same group were considered to not accurately reflect the effects of antibiotics.

A high interindividual variation of bacterial abundances was observed in all groups on day 0. The alpha diversity indices (Table S3), the ANOSIM of unweighted and weighted UniFrac distances (Table S4, Fig S1), and bacterial abundances (Table S1) did not differ significantly among groups on day 0.

### 2.1. Amoxicillin/clavulanic acid group

#### 2.1.A) Sequence analysis - alpha and beta diversity

The AMC group showed reduced evenness on the last day of treatment (day 20) compared to DOX and CON groups; this decrease approached but did not reach statistical significance (Shannon index, p = 0.061) (Table S3, Fig 4). A statistically significant difference in microbial community composition on the last day of treatment (day 20) was observed for AMC cats, compared to both DOX (ANOSIM R = 0.109, p = 0.011) and CON (ANOSIM R = 0.188, p = 0.001) cats based on unweighted analysis (Table S4, Fig 5). On days 60 and 300, there was a less distinct clustering of the microbiome in AMC cats compared to CON cats (based on decreasing ANOSIM effect size) as demonstrated by unweighted (ANOSIM day 60 R = 0.056, p = 0.075, ANOSIM day 300 R = 0.077, p = 0.058) and weighted distances (ANOSIM day 300 R = 0.057, p = 0.074), but this difference did not reach statistical significance (Fig S1).

**Fig 4.**
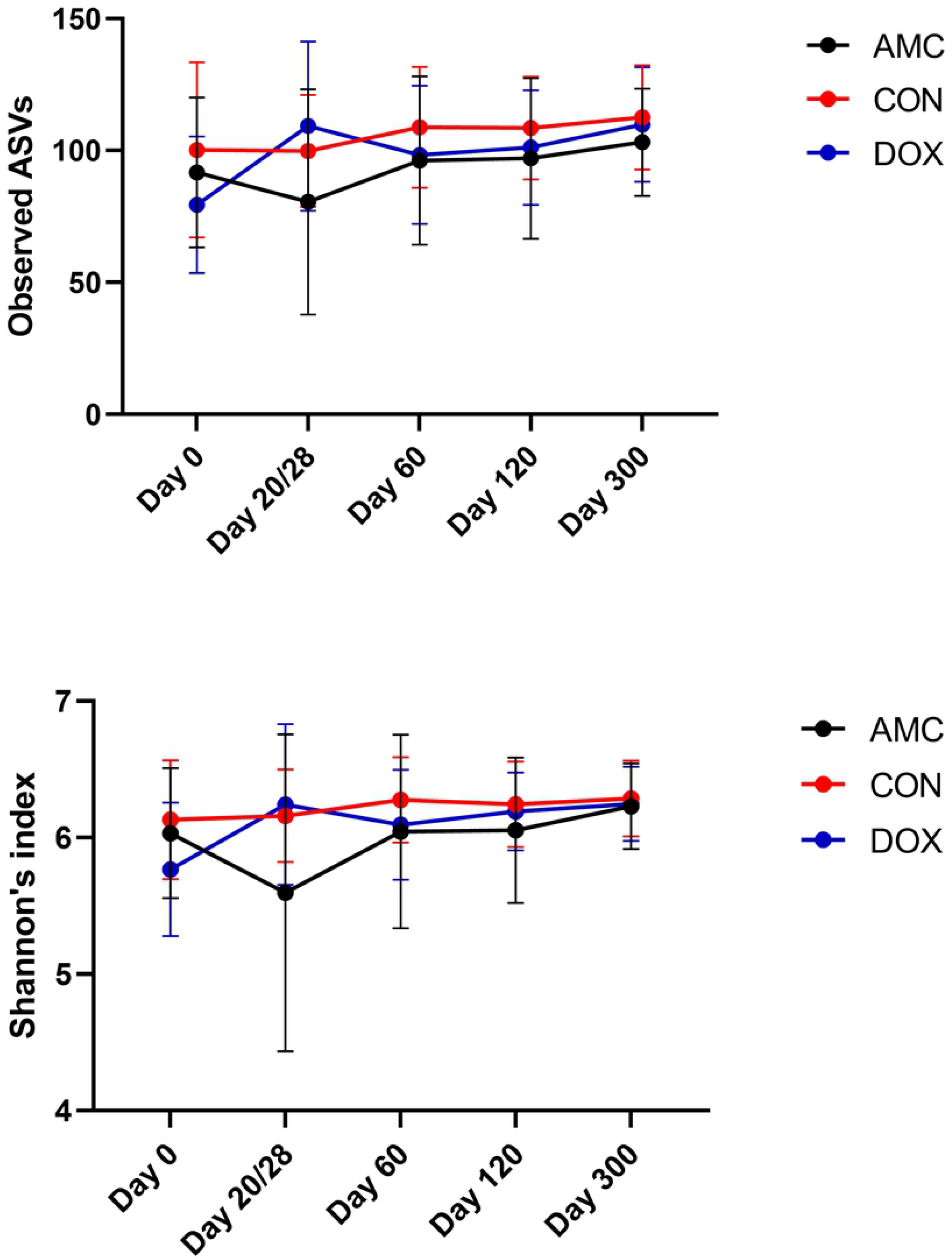
Alpha diversity differences between cats treated with amoxicillin/clavulanic acid (black), cats treated with doxycycline (blue), and healthy control cats (red). Means and standard deviations within each group are displayed.

**Fig 5.**
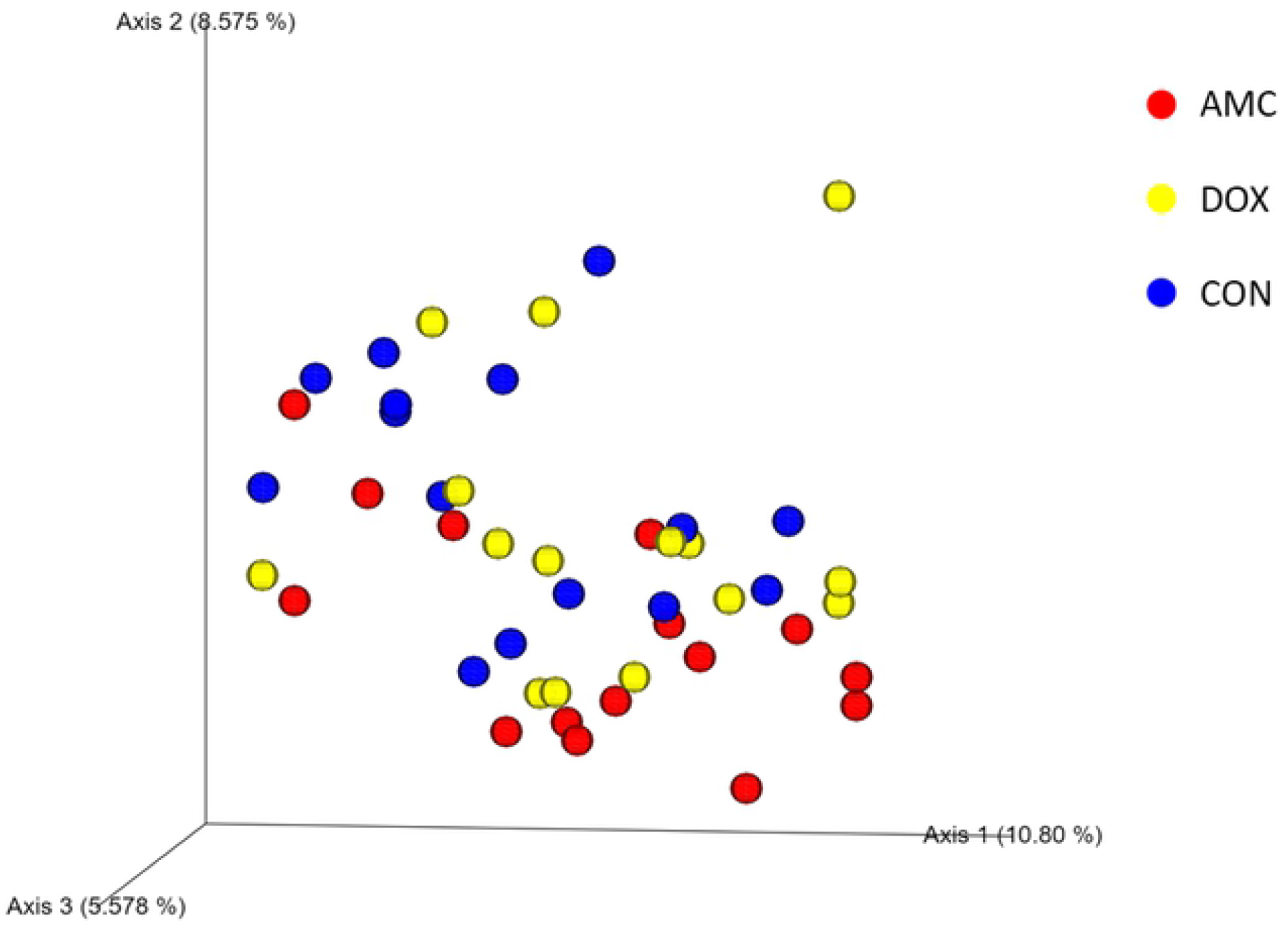
Principal Coordinate analysis (PCoA) plot of unweighted Unifrac distance in cats treated with amoxicillin clavulanic acid (red = AMC), cats treated with doxycycline (yellow = DOX), and control cats (blue = CON) on day 20/28.

#### 2.1.B) Sequence analysis - abundance of individual bacterial taxa

Amoxicillin/clavulanic acid had a significant impact on the GI microbiome. In fact, the normal age-related changes of the microbiome observed in CON cats were not observed in this group. Erysipelotrichi (p = 0.008), *Catenibacterium* spp. (p = 0.045), and unclassified Lachnospiraceae (p = 0.002) were detected in significantly lower abundances, whereas Enterobacteriales (p=0.010) was found in significantly higher abundances in feces from AMC cats compared to CON cats on the last day of treatment (day 20/28) (Figs 6 and 7). Three (day 120) and 9 months (day 300) after amoxicillin/clavulanic acid discontinuation, AMC cats harbored significantly higher abundances of unclassified *Collinsella* spp. compared to CON cats (Fig 7).

**Fig 6.**
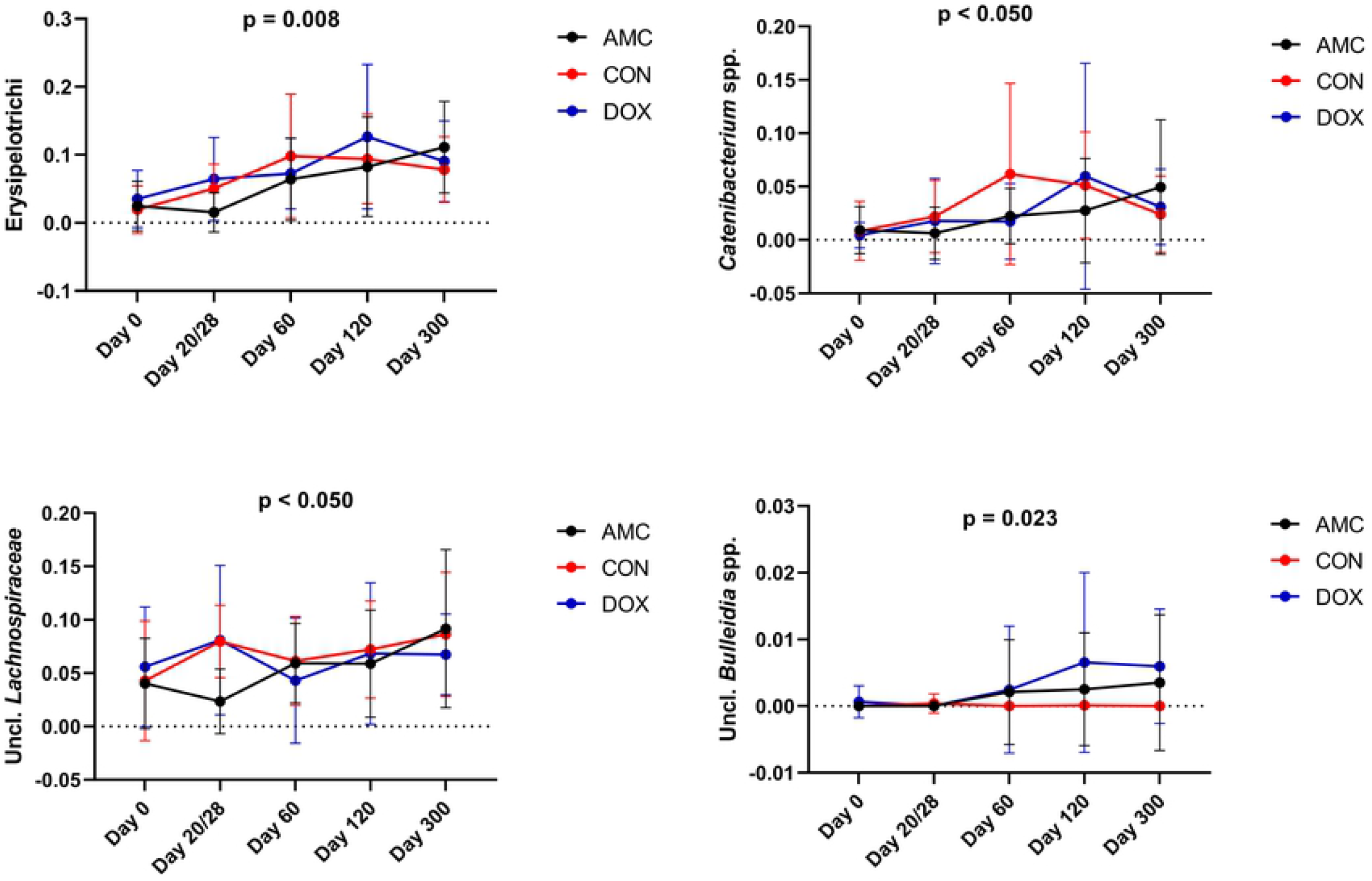
Bacterial groups that showed a significantly decreased abundance after antibiotic treatment (AMC and DOX group) compared to the control group (CON group). Means and standard deviations within each group are displayed.

**Fig 7.**
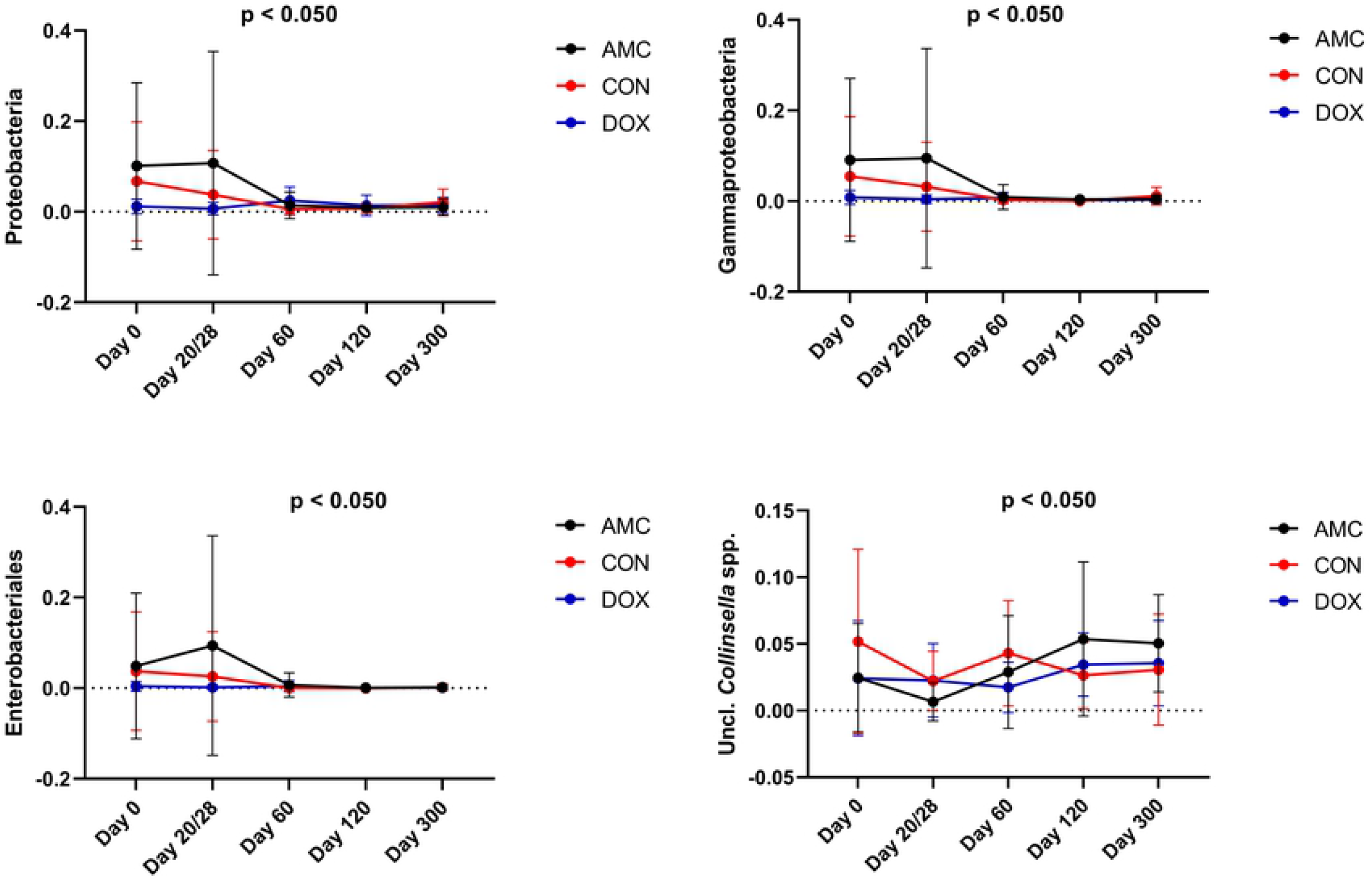
Bacterial groups that showed a significantly increased abundance after antibiotic treatment (AMC and DOX groups) compared to the control group (CON group). Means and standard deviations within each group are displayed.

Most of the differences in bacterial abundances between AMC and CON groups were found during treatment, (from 2 to 3 months of age), while after that period only minor changes were observed. In AMC cats, Gammaproteobacteria abundances remained the same during treatment (i.e., from 2 to 3 months) representing approximately 9% of total sequences, while in CON cats they decreased during the same period, representing 3% of total sequences (p = 0.009). At 1 month after antibiotic withdrawal (4 months of age), Gammaproteobacteria decreased to <1% in AMC cats (p = 0.030), reaching similar levels to those in CON cats at this age (Fig 7). Erysipelotrichi abundances represented 2.5% of the total sequences in AMC cats before treatment, and decreased to less than 2% after treatment, while in CON cats, Erysipelotrichi abundances increased at this age. On day 60, both groups harbored similar abundances of this bacterium (Fig 6).

#### 2.1.C) qPCR for selected bacterial groups

On the last day of treatment, lower total bacterial counts (p = 0.003) and higher abundances of *E. coli* (p = 0.002) were detected in the feces of AMC cats compared to CON cats (Fig 8).

**Fig 8.**
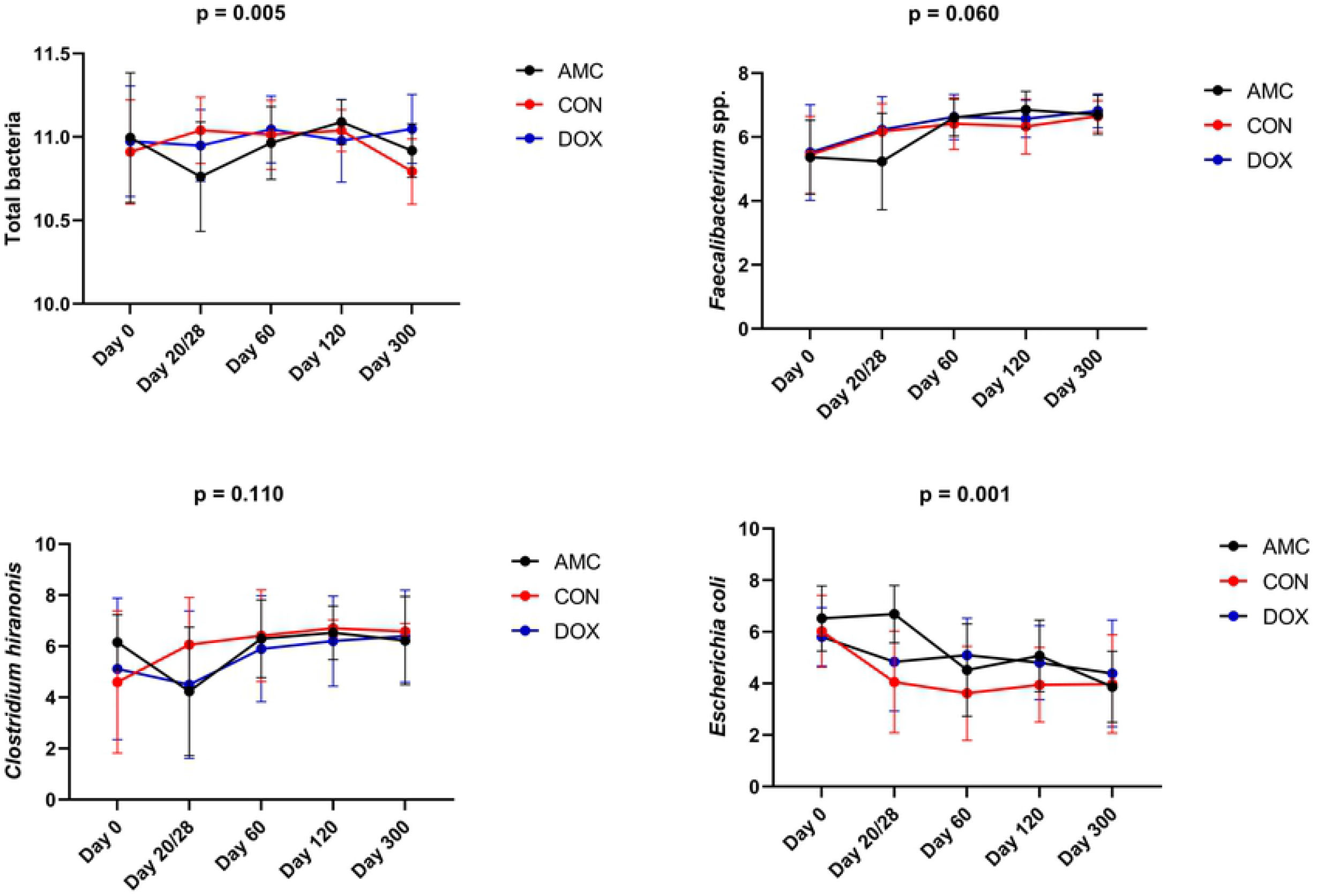
Fecal abundances of selected bacterial taxa among cats treated with amoxicillin/clavulanic acid (AMC), cats treated with doxycycline (DOX), and healthy cats (CON) analyzed with qPCR. Means and standard deviations within each group are displayed.

Bacterial abundances in the AMC group demonstrated a different pattern compared to the CON group. In the AMC group, *E. coli* abundances did not change between 2 to 3 months of age (i.e., during treatment), and then significantly decreased at 4 months of age (p = 0.012) (Fig 8).

### 2.2. Doxycycline group

#### 2.2.A) Sequence analysis – alpha and beta diversity

DOX cats had a significantly higher species richness (observed ASVs, p = 0.025; Chao1, p = 0.029) (Table S3, Fig 4) on the last day of treatment and a different clustering of the microbiome 1 month after treatment (day 60) compared to CON cats (ANOSIM R = 0.100, p = 0.021) (Table S4, Fig 9).

**Fig 9.**
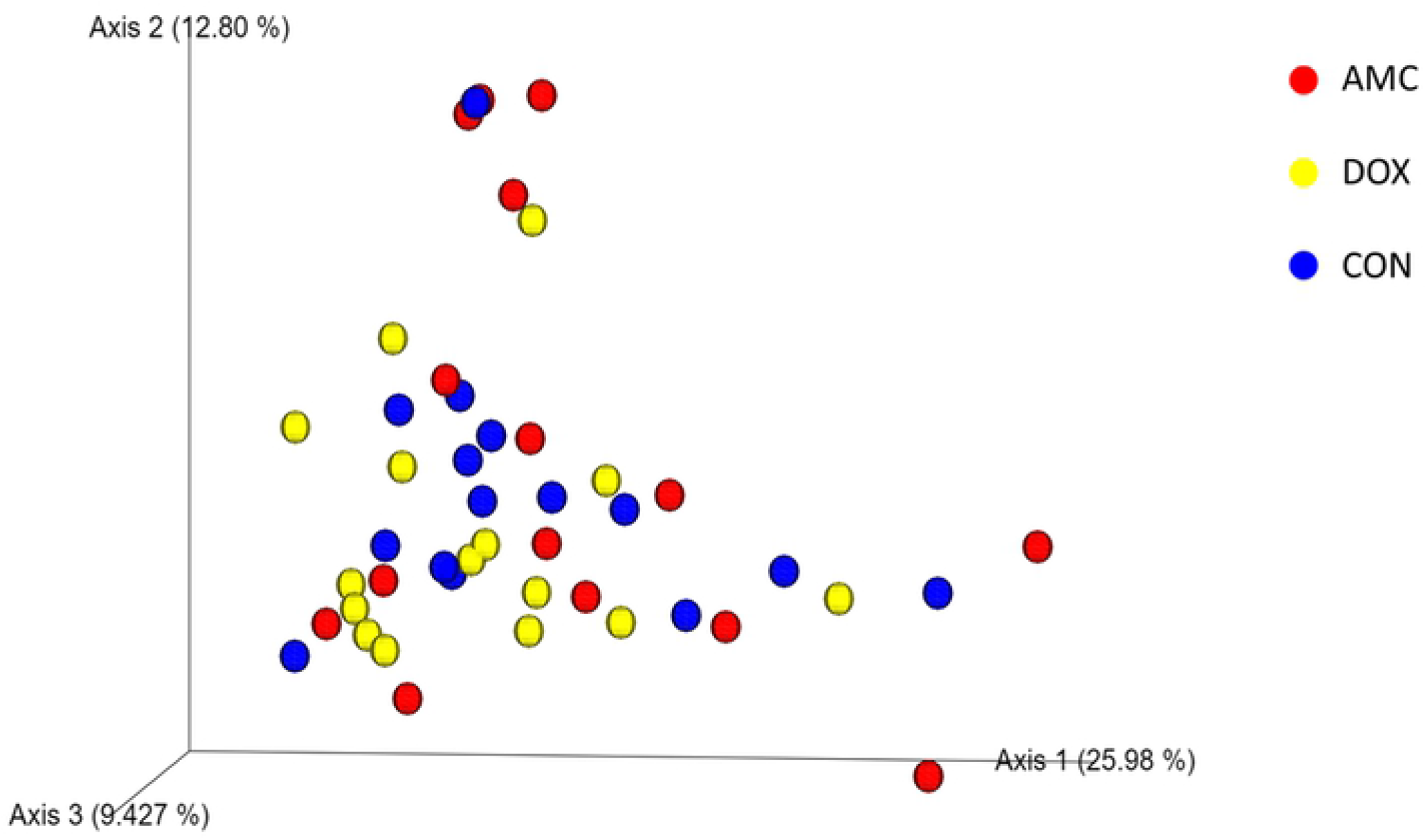
Principal Coordinate analysis (PCoA) plot of weighted Unifrac distances in cats treated with amoxicillin/clavulanic acid (red = AMC), cats treated with doxycycline (yellow = DOX), and control cats (blue = CON) on day 60.

#### 2.2.B) Sequence analysis – abundance of individual bacterial taxa

Doxycycline caused pronounced changes in the abundances of bacterial communities, but its effects appeared 1 month after its discontinuation. *Catenibacterim* spp., and unclassified *Lachnospiraceae* spp. (both p = 0.039) were detected at significantly lower abundances whereas Proteobacteria (p = 0.001) and Enterobacteriales (p = 0.018) at significantly higher abundances in the feces of DOX cats compared to CON cats on day 60 (Figs 6,7). The increase in the abundance of Proteobacteria persisted for 3 months after antibiotic withdrawal (p = 0.026). In addition, at 3 and 9 months after antibiotic withdrawal, the abundance of unclassified *Collinsella* spp. was significantly higher in cats of the DOX group compared to cats of the CON group (p = 0.025) (Fig 7). Unclassified *Bulleidia* spp. were detected at higher abundances (p = 0.023) in DOX cats 9 months after its discontinuation (Fig 6). Fig 10 shows a percentage plot of bacterial abundances at a class level among groups.

**Fig 10.**
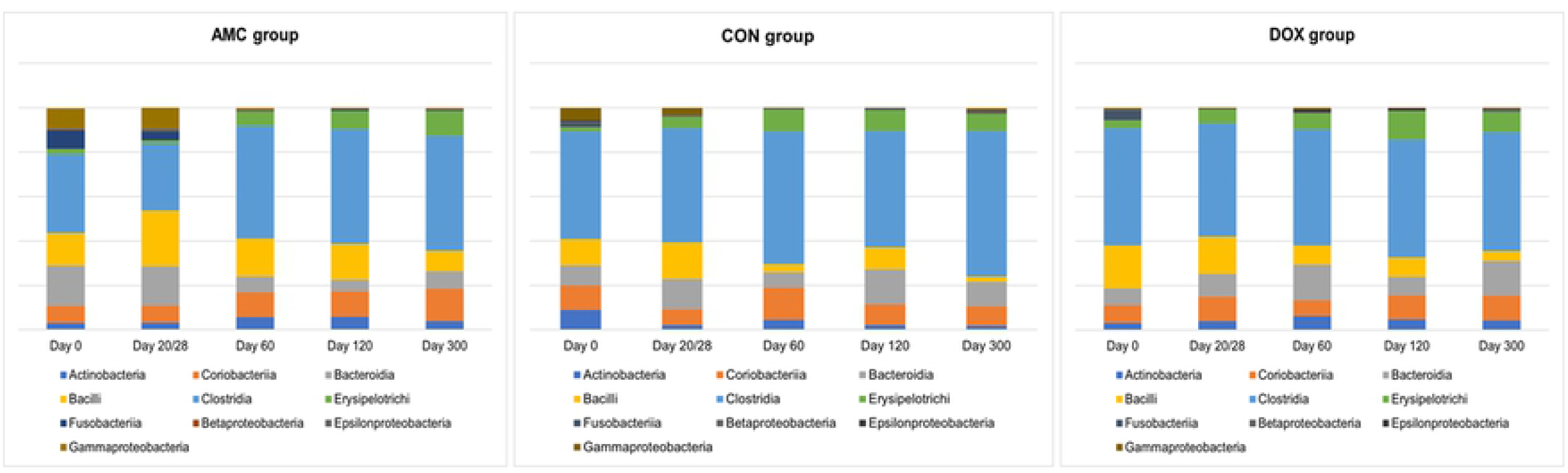
Relative abundance of bacterial taxa at a class level among groups.

#### 2.2.C) qPCR for selected bacterial groups

On day 60, higher *E. coli* abundances (p = 0.035) were found in DOX cats compared to CON cats (Fig 8).

## Discussion

Our goals were to describe the effects of treatment with amoxicillin/clavulanic acid or doxycycline on the GI microbiome of young cats and the microbial recovery after antibiotic exposure early in life. Our study showed substantial changes in the GI microbiome from 2 months until one year of age in cats, with antibiotics having a differential impact on the developing GI microbiome. Amoxicillin/clavulanic acid caused pronounced effects during treatment while the effects of doxycycline appeared 1 month after its withdrawal. Both antibiotics mainly affected members of Firmicutes and Proteobacteria and resulted in a delay in the developmental progression of the microbiome compared to the pattern of microbial changes observed over time in cats not treated with antibiotics.

Importantly, a high interindividual variation in bacterial abundances was observed in cats at 2 months of age (before exposure to antibiotics). In humans and dogs, during the phase of microbiota maturation, high-interindividual differences in bacterial abundances occur [45-47], therefore the large variation observed in our study likely represents an immature microbiome in cats at 2 months of age. In addition, the largest shifts in the GI microbiota in the control cats occurred during the age of 2 to 6-months suggesting that the normal GI microbiome evolves in kittens and reaches maturity around the age of 6 months. Although conflicting evidence exists about whether the microbiome reaches an adult-like state at the end of the weaning period in dogs and cats [8, 48, 49], in a previous canine study, 2-month-old puppies still harbored a significantly different microbiome compared to adult dogs [45].

In adult humans, the abundances of approximately 70% of the GI bacterial members is relatively stable for at least 12 months [8]. Therefore, in contrast to adult cats, the duration of antibiotic effects on the developing GI microbiome could only be investigated by evaluating a control group to adjust for age-related changes. The fact that there was a large variation in microbial community composition at baseline among cats likely led to unique responses to antibiotics. The microbiome is considered as unique as an individual’s fingerprint [50], and during the maturation period, unpredictable shifts could occur that have not been adequately described in cats. Despite the high variability, the core bacterial taxa in cats of our study were Firmicutes and Actinobacteria from 2 months until 1 year of age. This is in agreement with previous studies investigating the effects of dietary nutrient composition [48, 51, 52], sex, and sexual status [49] on the fecal microbiome of young cats.

Current knowledge suggests that the first microbes colonizing the GI tract are mainly facultative anaerobic bacteria that reduce oxygen concentrations in the gut and allow for successful colonization of the obligative anaerobic bacteria [53]. The phylum Proteobacteria, which is comprised by facultative and obligative anaerobic bacteria, is among the first colonizers of the GI tract in humans [53, 54]. At the weaning period and after the introduction of a solid diet (i.e., around 5-6 months of age), the abundance of Proteobacteria gradually decreases [55]. Our finding of an age-dependent decrease in bacterial taxa belonging to Proteobacteria (i.e., Enterobacteriales, Escherichia coli) observed between 2 to 4 months of age in control cats in this study is in agreement with these data in humans. In addition, a concurrent increase in the abundance of taxa belonging to Firmicutes (i.e., Erysipelotrichales) occurred in the same group during the same period, which has also been reported by another study in cats of a similar age and reflects the introduction of dietary macronutrients that are utilized by these bacteria [51].

Amoxicillin is a semisynthetic penicillin that is active against some non-beta-lactamase producing gram-positive bacteria and few gram-negative bacteria. The addition of a beta lactamase inhibitor, such as clavulanic acid, increases the spectrum of activity of amoxicillin [56]. Doxycycline belongs to tetracyclines, a class of bacteriostatic antibiotics with broad spectrum activity against bacteria, rickettsiae, and protozoal organisms. Tetracyclines are also known for their anti-inflammatory properties, which seem to contribute to their therapeutic efficacy [57]. These antibiotics constitute two of the most commonly prescribed antibiotics in young cats.

Treatment with amoxicillin/clavulanic acid led to a trend in reduced species richness and evenness, although this varied among cats and it did not reach statistical significance. Similarly, in one study in adult laboratory cats, amoxicillin/clavulanic acid for 7 days reduced the number of different species observed and this effect persisted for 7 days after discontinuation of the antibiotic [30]. In our study, species richness indices were indistinguishable from untreated cats by 1 month after discontinuation of amoxicillin/clavulanic acid.

Doxycycline had a different effect on the GI microbiome. Increased abundance of different species were observed by the end of treatment. Antibiotics, including doxycycline and amoxicillin most commonly either decrease [58-62], or do not have any effect on species richness [63]. Only few studies have reported an increase in species richness indices [64, 65]. In our study, doxycycline had no effect on bacterial abundances and community composition on the last day of the treatment period (day 28). Alternatively, the lack of an effect of doxycycline on bacterial genera that would be expected to decrease as shown in control cats, could be responsible for the observed increased species richness in doxycycline-treated cats. The bloom of these genera might be attributed either to resistance to tetracyclines or to the concurrent decrease of some bacteria that produce antimicrobial peptides thus allowing members of these genera to remain at increased levels [66].

Microbial community composition was distinct in cats treated with amoxicillin/clavulanic acid and indistinguishable in cats treated with doxycycline compared to control cats on the last day of treatment. Interestingly, the effect of doxycycline was not evident until 1 month after its discontinuation of the drug.

Similar results have been described in a single study in mice, where the most profound changes in microbial community composition started 1 month after doxycycline discontinuation [67]. In addition, in our study, a trend for significant differences in microbial community composition were observed in amoxicillin/clavulanic acid-treated cats 3 and 9 months after antibiotic withdrawal.

Contradictory findings exist in the literature with humans, laboratory animals and in vitro studies reporting high interindividual effects [68], no effects [69, 70], only short-term effects [62, 65], or both short- and long-term effects on microbial composition [60, 71, 72] after administration of amoxicillin with or without clavulanic acid. In a study in rats, a 7-day course of amoxicillin during the weaning period caused transient alterations in microbial composition that resolved by 20 days after its discontinuation [61]. In another study in infants, a 5- to 8-day course of amoxicillin caused long-term changes in microbial composition that persisted for 6 months after treatment withdrawal [72].

While the total abundance of the phylum Firmicutes was not significantly altered, certain bacterial members of this phylum showed significant shifts in response to antibiotics. Amoxicillin/clavulanic acid and doxycycline administration caused a transient decrease of the abundance of the order Erysipelotrichales and its sub-groups Erysipelotrichaceae and *Catenibacterium* spp. The family Erysipelotrichaceae contains bile salt hydrolase (BSH) genes, and this enzyme is responsible for the deconjugation of primary bile acids [73, 74]. Thus, the decrease observed could potentially lead to increased concentrations of deconjugated primary bile acids in the gut. In addition to potential bile acid dysmetabolism in cats treated with antibiotics, one of the main converters of primary bile acids into secondary bile acids in dogs and cats is *Clostridium hiranonis*, which showed a decreased abundance in response to both antibiotics in our study, although this change did not reach statistical significance for either treatment [75]. Families belonging to Clostridiales were affected by antibiotics with a significant decrease in unclassified Lachnospiraceae. The family Lachnospiraceae was the predominant family present at all time points in all groups.

Members of this family ferment carbohydrates leading to the production of butyrate [76]. Butyrate is one of the main short chain fatty acids (SCFAs) in the gut and has anti-inflammatory properties, is a major energy source for colonocytes, and its absence causes autophagy of epithelial intestinal cells in germ-free mice [77, 78]. As a result, SCFAs might be another main metabolic class influenced by antibiotic treatment. A more comprehensive picture of the antibiotic effects on the GI microbiome could therefore be obtained by applying other “omics” approaches including metabolomic analysis leading to a better understanding of the metabolic pathways affected by antibiotics.

Among Actinobacteria, the abundance of unclassified *Collinsella* spp. was higher in both antibiotic-treated groups than in controls at 3 months after discontinuation of treatment. This effect persisted in the amoxicillin-clavulanic acid group for 9 months. Early colonization with *Collinsella* spp. within the first 6 months of life is associated with increased adiposity in humans, [55] and also increased *Collinsella* spp. abundances have been reported in cats with diarrhea [79, 80].

Based both on sequencing and qPCR analysis, bacterial taxa belonging to Proteobacteria (Gammaproteobacteria, order Enterobacteriales, family Enterobacteriaceae, *Escherichia coli*) were found at significantly higher abundances on the last day of treatment (20 days) for amoxicillin/clavulanic acid and at 3 months after discontinuation of doxycycline before decreasing to similar abundances to that of control cats. The family Enterobacteriaceae is the most common microbial member that increases in abundance after antibiotic treatment in humans regardless of the antibiotic class [81]. In dogs, metronidazole [82] and amoxicillin [83], but not tylosin [84, 85], are reported to increase the abundance of Enterobacteriaceae. In cats, this effect has been observed for amoxicillin [30] and clindamycin [31, 32] with the latter leading to a 2-months persistent increase in Enterobacteriaceae [32]. The phylum Proteobacteria encompasses some of the most well-known pathogens [54] and members of this phylum are commonly increased in dogs [86-91] and cats with GI disease [79, 80, 92-94], as well as during consumption of high-protein, canned and raw diets [48, 51, 52, 95]. Both antibiotic treated groups had higher fecal scores during treatment compared to healthy cats, therefore episodes of diarrhea may be associated with increased abundances of Proteobacteria members.

Previous studies in humans have shown that antibiotics delay the developmental progression of the microbiome into an adult-like state [21, 22]. In agreement with these findings and compared to untreated cats of our study, a delay in maturation was observed in both antibiotic-treated groups. This delay was characterized by reduced abundances of taxa belonging to Firmicutes and increased abundances of taxa belonging to Proteobacteria. The most profound delay occurred between 2 to 3 months of age in the amoxicillin/clavulanic acid-treated cats and between 3 to 6 months of age in the doxycycline-treated cats.

Our study had some limitations. All cats were stray at study initiation; thus, their exact date of birth was unknown and slight differences in the enrollment age might have influenced the microbiota composition. Some cats were malnourished, and malnourishment has been associated with a persistently immature microbiome in children [96]. In addition, some cats were found at a very young age and required formula feeding, which in children is also reported to impact microbiome colonization compared to breastfeeding [97]. The maternal diet of cats also has an impact on the microbiome of the offspring until its 17th week of age [98] and in our study the maternal dietary status was unknown. Although the above factors have been investigated in humans, no studies regarding their impact on the feline microbiota exist. Finally, cats treated with doxycycline had a significantly higher fecal scores at baseline, which might also have influenced the abundance of some bacterial taxa.

## Conclusion

Overall, our results indicate that the GI microbiome of cats changes after 2 months of age and reaches an adult-like state around 6 months of age. Amoxicillin/clavulanic acid and doxycycline treatment early in life significantly affected the developing microbiome richness and composition in cats. The abundance of members of Firmicutes decreased and that of members of Proteobacteria increased after 20 days of amoxicillin/clavulanic acid treatment and 1 month after a 28-day course of doxycycline. Only minor changes were observed 9 months after amoxicillin/clavulanic acid or doxycycline discontinuation with an increase in the abundance of unclassified *Collinsella* spp. and unclassified *Bulleidia* spp., respectively. Our results suggest that doxycycline had a delayed impact whereas amoxicillin/clavulanic acid had a more immediate impact on bacterial community composition and only minor changes persisted 9 months after discontinuation of either antibiotic. Future studies utilizing additional approaches to gain a better understanding of the microbial functional changes caused by antibiotics would be useful.

## Acknowledgments

Preliminary results were presented at the: a) 29^th^ European College of Veterinary Internal Medicine Congress, Milan, Italy, September 19^th^ to 21^st^ 2019; b) the 38^th^Forum of the American College of Veterinary Internal Medicine (ACVIM), June 10^th^ to13^th^ 2020; and c) the 39^th^ Forum of the American College of Veterinary Internal Medicine (ACVIM), June 9^th^ to12^th^ 2021.

The authors would like to thank Gerolymatos International S.A. for providing products for antiparasitic treatment (Broadline) and vaccines (Purevax RCPh, Purevax Rabies) for the cats in this study.

## Supporting information

**S1 Fig. Beta diversity indices among groups**. A) Principal Coordinate Analysis of unweighted UniFrac distances of 16S rRNA genes representing the difference in microbial communities among cats treated with amoxicillin clavulanic acid (blue circles), cats treated with doxycycline (yellow circles), and healthy control cats (red circles) on days 20/28 (last day of treatment), 60, 120, and 300. B) Principal Coordinate Analysis of weighted UniFrac distances of 16S rRNA genes representing the difference in microbial communities among cats treated with amoxicillin clavulanic acid (blue circles), cats treated with doxycycline (yellow circles), and healthy control cats (red circles) on days 20/28 (last day of treatment), 60, 120, and 300.

**S1 Table. Summary statistics of sequencing data describing the mean percent and standard deviation of sequences belonging to antibiotic-treated (AMC and DOX groups) and healthy (CON group) cats**.

**S2 Table. Summary statistics of qPCR data describing the mean log abundance and standard deviation of bacterial groups belonging to antibiotic-treated (AMC and DOX groups) and healthy (CON group) cats**.

**S3 Table. Alpha diversity metrics (mean ± standard deviation). CON, healthy cats that did not receive antibiotics; AMC, cats treated with amoxicillin/clavulanic acid for 20 days; DOX, cats treated with doxycycline for 28 days**.

**S4 Table: Beta diversity differences. CON, healthy cats that did not receive antibiotics; AMC, cats treated with amoxicillin/clavulanic acid for 20 days; DOX, cats treated with doxycycline for 28 days**.

